# Modeling neural activity in neurodegenerative diseases through a neural field model with variable density of neurons

**DOI:** 10.1101/2022.08.23.504980

**Authors:** Ronaldo García Reyes, Eduardo Martinez Montes

**Author notes:** **Mail:**.

## Abstract

In recent years, a vertiginous advance has occurred within the Neural Field Theory with the development of the so-called Next Generation Neural Field models. Unlike the phenomenological models, these models manage to describe neuronal activity, macroscopically, from the thermodynamic limit of microscopic laws under the assumption of a homogeneous density of neurons. The study of neural activity during neurodegenerative processes associated to Alzheimer’s, Parkinson’s or Glioblastomas, should include a variable density of neurons. In this work, we propose an update of the Next Generation Neural Field model, extracted from the thermodynamic limit of the quadratic integration-and-fire model with realistic synaptic coupling and a variable density of neurons at the microscopic level. The thermodynamic limit of the system will allow us to study the patterns of synchronized neural activity that appear as the result of different spatial distribution of neurodegeneration. In particular, we demonstrate that during neurodegenerative processes, the relationship established between the thermodynamic states of the Neural Field and the Kuramoto order parameter (Measure of Neural Synchronization) differs from the classic results of the Next Generation Neural Field literature. Instead, the variation in neuron density directly modifies the Kuramoto order parameter. This might help us explain the diverse patterns of activity that can be found in different neurodegenerative processes and that could become experimental biomarkers of such pathologies.

## 1. Introduction

Neurodegenerative diseases (NDDs) refer to a wide group of diseases of the nervous system of diverse origin with various neuropathological and clinical manifestations. Patients with these diseases may experience alterations in their cognitive functions or movements. Despite the various clinical manifestations or origin, NDDs are characterized by a progressive process of neural degeneration and death of neurons in the nervous system. Examples of neurodegenerative diseases are Alzheimer’s (AD), Parkinson’s (PD), Huntington’s (HD) and other conditions that lead to neurodegeneration (i.e., the death of neurons) such as Glioblastoma multiforme (GBM). The different neuropathological and clinical manifestations depend on the areas of the nervous system where, neuronal death takes place [You09, Mat00].

Previous studies have established that early detection of neurodegeneration is good [PNK^+^13]. Therefore, the development of novel diagnostic techniques of NDDs in the initial stages is currently a critically important goal. For this purpose, it is necessary to identify biomarkers with predictive and minimally invasive capacity. In articles such as Nimmrich *et al*. [NDA15], the authors have identified biomarkers extracted from the Electroencephalogram (EEG), Functional Magnetic Resonance Imaging (fMRI) among others that seek to diagnose NDDs based on their relationship with neuronal activity. In Aron *et al*. [AY16] and Iram *et al*. [AJIV^+^15, IVQ16] the authors propose to use EEG signals to estimate neural synchronization as a biomarker in the early diagnosis of AD. In Asmarian *et al*. [ARE^+^21] the authors propose a rating scale method for PD based on the strength of synchronization between dynamics of strides. On the other hand, investigations have emphasized a relationship between large-scale synchronization of oscillations in the *β* - frequency band and akinesia^1^ [Bro03, SG05]. Finally in the article Uhlhaas *et al*. [US06], the authors carry out a general bibliographic review of the literature, they show results that suggest a close correlation between abnormalities in neuronal synchronization and cognitive dysfunctions such as NDDs.

These studies and others have shown that neurodegeneration induces changes in functional synchronization of neural activity in initial stages, which suggests that the study of synchronization patterns reflects the presence of neurodegeneration. This might be simply explained by the death of neurons that belong to the same network (decreasing the effective number of neurons that can be synchronized) or by the impairment of nervous fibers connecting the neurons in the network and disrupting the functional synchronization.

Despite the importance of the relationship between neuronal activity and NDDs, most of the articles in the literature limit themselves to studying this relationship from the phenomenological point of view. These phenomenological descriptions do not allow us to answer all the questions, since it does not maintain a direct connection with the microscopic reality of the NDDs. In other words, this means that phenomenological models are unable to establish why biomarkers taken from the EEG, such as synchronization, are adequate for the diagnosis of NDDs. We propose that this type of question can be answered if we model this relationship using a bottom-up approach of the Neural Field Theory (NFT) or a version of the New Generation Neural Field models (NGNF).

Neural Field Theory (NFT), as defined by Coombes *et al*. [Coo06], is the branch of Theoretical Neuroscience that aims to describe neural activity within a space-time region. The mathematical models that make up this theory can be classified according to the bottom-up or top-down approach. The class of top-down models is made up of phenomenological equations extracted from the emergent behavior. Within this class we can include the classic Wilson-Cowan models, the Amari model and the Jensen-Rit model as presented in Cook *et al*. [CPWT21]. One of the main disadvantages of this class of models is that it does not maintain a strong connection with the microscopic reality of the problem. The class of bottom-up models is made up of models that intend to describe the response of individual neurons or neuronal masses at the microscopic or mesoscopic level, and then integrate them to obtain a mean-field description. Achieving the integration of neural activity in the bottom-up approach is especially difficult due to the high complexity of the brain tissue.

As defined by Byrne *et al*. [BAC19] the models of NFT that belong to the bottom-up approach that incorporate within-population synchrony and a realistic form of synaptic coupling, make up the NGNF models. In the literature we can find several examples of NGNF models, most of them are obtained from the thermodynamic limit of the quadratic integrate- and-fire model (QIF) with realistic synaptic coupling that describes the high frequency type I cortical neuron. The exact thermodynamic limit for a version of the QIF model is determined by applying the Otto-Antonsen Ansatz (OA) over the equivalent *θ*-model [BRNC22] or using Lorentzian Ansatz (LA) [MPR15]. This allows the modeling of macroscopic neural activity within a region Ω ⊂ ℝ^2^ by using the spatial-temporal distribution of 4 thermodynamic state variables: the firing rate *R*, the average potential *V*, the synaptic activity *U* and the spatial-temporal drive Ψ. These variables can be obtained as the continuum thermodynamic limit of the QIF models of neurons in Ω region, then is possible to compute the local synchronization^2^ with the Kuramoto order parameter *Z* from the thermodynamic state (*R,V,U*, Ψ) by the following relation [BRNC22, MPR15]

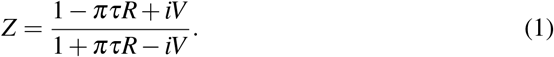

It is known that neuronal death in a brain region is a pathological characteristic for different NDDs Chi *et al*. [CCS18]. For example, PD is due to the loss of dopaminergic neurons in the substantia nigra [CA05b] while AD is due the loss of basal forebrain cholinergic neurons [AKBQ02, GA98]. In the spatial-temporal framework of NFT neuronal death is described from a variable neuron density *ρ*. Previous studies have already used neuronal density as an anatomical feature of NDDs. For example, in Fearnley *et al*. [FL91] the authors report that neurodegenerative process in PD follows a negative exponential course, result that was confirmed by new data in [HSC^+^05, ZRPS06, CCL^+^00, CL05]. Based on these results we will hypothesize that the space-time evolution of the neuronal density *ρ* i.e., the neurodegenerative process studied evolves according to an exponential decay model (subsection 2.1.2).

Our main objective during this article is to develop a mathematical model to describe neural activity in the presence of neurodegenerative processes. For this, we decided to work on the theoretical framework of NGNF models, since this allows us to introduce the effects of neurodegeneration on individual neurons by an update of microscopic QIF model, then integrate their responses through the thermodynamic limit to finally find the macroscopic description. From the above we can conclude that to model neural activity during NDDs within the framework of NGNF models we need to couple a variable related to the density of neurons in the field equations. To our knowledge, there is no extension in the literature of the NGNF models for a variable density of neurons, nor is there a study on the implications of neurodegeneration on the synchrony of neurons using a bottom - up approach.

In section 2.2 we will deduce the update NGNF model Eqs. (4) for a variable density of neurons, we show that in a presence of a neurodegenerative processes the macroscopic state its described by fifth variables (*R,V,U, ψ, λ*), where *λ* its directly related with neuron density (subsection 2.1.2) and *ψ* it is the analog of the spatial - temporal drive Ψ for neurodegenerative processes (subsection 2.1.3). Later, during the exposition of the theoretical results (subsection 3.1), we will analyze the relationship that is established between the macroscopic state and the Kuramoto order parameter. We will show that the expression (1) is no longer valid for neurodegenerative processes, instead the appropriate expression is

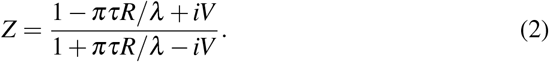

Form the Kuramoto order parameter we will define and study general properties of the *K* - point synchronization 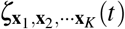 that it is a measure a synchronization between *K* point inside Ω (subsection 3.1). Then we study the consistency of our model evaluating the results of the numerical simulations (subsection 3.2).

## 2. Mathematical model

### 2.1 Model hypothesis

We have divided the exposition of the hypothesis of the model into three parts. During the first one, we will present the classic NGNF model, which we will assume describes neuronal activity in the absence of neurodegenerative processes i.e health and constant density of neurons *ρ*_*H*_. During the second part we will introduce a new variable that is related to the density of neurons *ρ* and can be interpreted as the *probability* that a neuron at a given point is alive and we will denote it by *λ*. We will use *λ* to describe the progression of neurodegenerative disease instead of neuron density *ρ*. In the last part we will describe how structural connectivity is affected by the neurodegenerative process described by *λ*. We are also going to introduce a new variable, the conditioned spatial - temporal drive, which we will denote by *ψ*, this new variable will allow us to describe the evolution of the system in a more elegant and effortless way.

#### 2.2.1 NGNF for non neurodegenerative processes

The main hypothesis of the classical NGNF model [BRNC22], is that in absence of neurodegenerative processes i.e. health and constant density of neurons *ρ*_*H*_, the neural activity within each neighborhood *ℬ*_*ε*_ (**x**) ^3^ of Ω can be described from a neural network of *N* QIF neurons with realistic synaptic connections. To simplify the notation we are going to denote the voltage of the neuron in **x**_*i*_ ∈ *ℬ*_*ε*_ (**x**) simply as *v*_*i*_ instead of *v*(**x**_*i*_, *t*) and for the input values of the other variables of the QIF model as Ψ we will denote Ψ = Ψ(**x**, *t*).

Assuming that the voltage *v*_*i*_ satisfies the QIF model with both gap-junction and synaptic coupling we have inside all *ℬ*_*ε*_ (**x**) of Ω

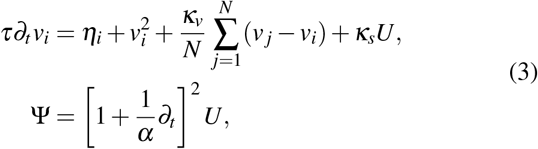

where *i* = 1, · · ·, *N* denote the neurons inside *ℬ*_*ε*_ (**x**), *v*_*i*_ is the voltage of the neuron *i* subject to reset *v*_*i*_ → *v*_*r*_ whenever *v*_*i*_ ≥ *v*_*th*_, *τ* is the membrane time constant, *k*_*v*_ and *k*_*s*_ are the strengths of gap-junction and synaptic coupling respectively, *η*_*i*_ is the random background input of the neuron *i* with Lorentzian distribution of *n*_0_ mean and *γ* half width^4^, *α*^−1^ is the time-to-peak of the synapse. *U* is the synaptic activity and Ψ the spatial-temporal drive that measures nonlocal interaction between all neighborhood of Ω with *ℬ*_*ε*_ (**x**).

From Eq. (3) we can obtain the macroscopic description of the problem by finding the continuous thermodynamic limit, which consists in considering that the number of neurons *N* that belongs to the neural network within each neighborhood *ℬ*_*ε*_ tends to infinity (*N* → +∞) while the size of each neighborhood becomes insignificant (*ε* → 0). Using the LA, the thermodynamic limit of the Eq. (3) can be found for a constant density of neurons as

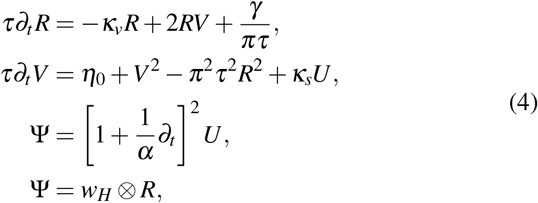

where the symbol ⊗ is used to describe spatial interaction within the neural field model ^5^, *w*_*H*_ is the structural connectivity in the absence of neurodegenerative processes and it is assumed that it satisfies the exponential decay model *w*_*H*_(y) = exp(−σ∥**y**∥)/(2*σ*). *ν* represent the speed of an action potential that takes values between 5 - 10 m/s in humans. For us, this model will be the one that models neuronal activity during non-neurodegenerative processes, that is, a healthy and constant density of neurons *ρ*_*H*_. We can appreciate a simulation of this model in Fig. 1.

**Figure 1.**
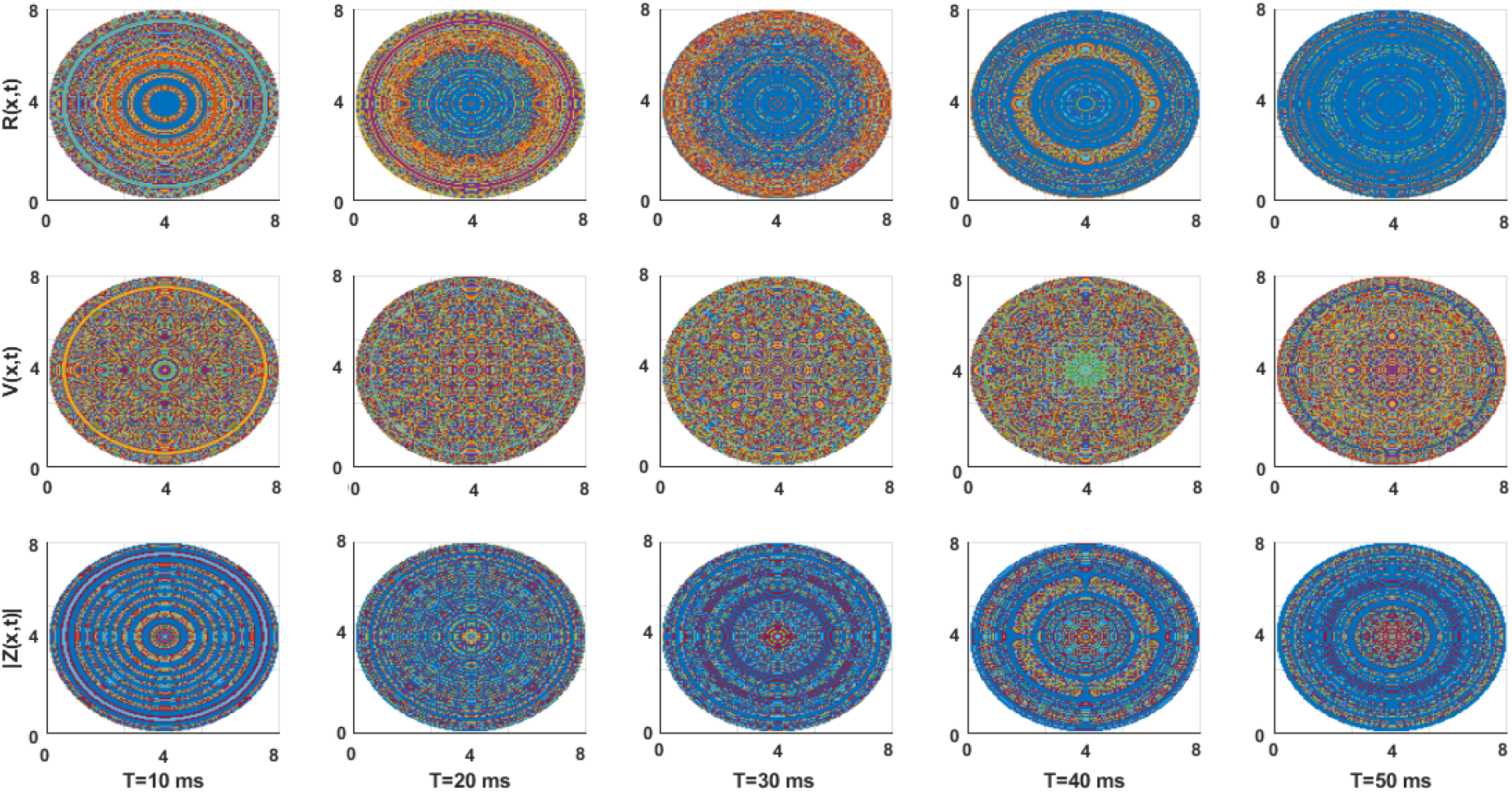
Simulation of the NGNF for a non neurodegenerative processes (*λ* = 1) inside a circumference Ω (with ratio *R* = 8). Values of the other constants: *τ* = 10, *α* = 3, *ν* = 10, *β* = 0, *v*_0_ = 0, *κ*_*v*_ = 0.9, *κ*_*s*_ = 5, *η*_0_ = 0.3, *γ* = 0.5, *σ* = 1.

#### 2.1.2 Dynamics of the neuron density

As we explained during the Introduction 1 several authors have considered describing the progress of a neurodegenerative process based on the density of neurons *ρ*. They further propose that the neuron density *ρ* must follow an exponential decay during NDDs, in mathematical terms this translates to

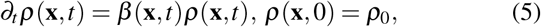

for all **x** in Ω, where *β* is a function that varies depending on the neurodegenerative process that we study and *ρ*_0_ is the initial density of neurons at the point **x**. In the case of neurodegenerative processes the density of neurons will be lower than the density of neurons in the absence of NDDs i.e. the health neuron density. This means that *ρ*(**x**, *t*) ≤ *ρ*_*H*_.

Using the definition of neuron density we can compute the number of living neurons at any moment *N*_*ε*_ (*t*) and maximum number of living neurons possible *N*_*H*_ inside all ball *ℬ*_*ε*_ (**x**) ∈ Ω. Then the probability of finding living neurons within the ball *ℬ*_*ε*_ (**x**) is

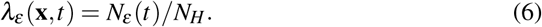

Since the neighborhood dimensions are negligible at the continuous limit (*ε* → 0), we can neglect the probability dependence of *ε*. Using the definition of *λ* it is possible to deduce from Eq. (5) the following equation (Appendix 5.1)

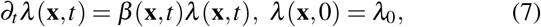

where *λ*_0_ = *ρ*_0_*/ρ*_*H*_. Since the new variable introduced to describe the degree of neurodegeneration is a probability, it is a normalized measure that takes values from 0 (absolute neurodegeneration) to 1 (absence of neurodegeneration).

#### 2.1.3 Conditioned spatio - temporal drive *ψ*

Structural connectivity is a measure of the anatomical connections between different areas of the brain, and it has been observed that it undergoes relevant changes during neurodegenerative processes, therefore we can find several articles in the literature that propose use brain connectivity as biomarker for NDDs, functional connectivity [PFVDH^+^14a].

In Eqs. (4) of the spatial-temporal drive Ψ, we can appreciate a term related to the structural connectivity *w* = *w*(**x, x**′) and it can be interpreted as the probability that an action potential propagates between the points **x** and **x**′ of Ω [CPWT21]. To model the structural connectivity function *w* during neurodegenerative processes we must consider the neurodegeneration within Ω described by *λ*. Therefore, we will assume that the integrity of the transmission of a nerve stimulus between two neurons depends only on the probability of find this neuron in the live state, which in terms of the probabilistic interpretation of the structural connectivity function *w* translates into the following relationship

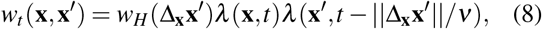

where *w*_*H*_ is the structural connectivity in the absence of a neurodegenerative process. From Eq. (8) we can express the spatial-temporal drive as

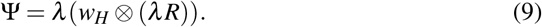

For convenience will denote *ψ* = *w*_*H*_ ⊗(*λ R*) and we will call her conditioned spatial - temporal drive. The main reason for the name is that *ψ* represents the value of Ψ conditioned to the fact that the post synaptic neuron is alive or *λ* (**x**, *t*) = 1.

We can exploit the convolutional structure and the spatial kernel shape *w*_*H*_ to reduce this integral to a Partial Derivative Equation (PDE) and get an equivalent to the Brain Wave Equation during neurodegenerative processes [Coo06].

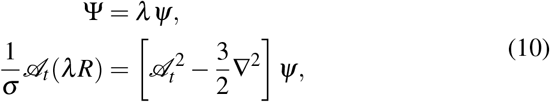

where *𝒜*_*t*_ = 1*/σ* + *∂*_*t*_*/ν* (Appendix 5.2).

### 2.2 NGNF model for neurodegenerative process

During the subsection 2.1.1 we explained that Eqs. (3) constitutes the microscopic description of neural activity in the absence of NDDs, therefore, it will be our starting point to develop our update NGNF model. To obtain a realistic microscopic description for neurodegenerative processes, we must extend the QIF model given by Eq. (3) considering the probability of finding a live neuron *λ* and his dynamic Eq. (7). For this purpose, it is convenient to express the Eq. (3) for the voltage *v*_*i*_ within *ℬ*_*ε*_ (**x**) in integral form as

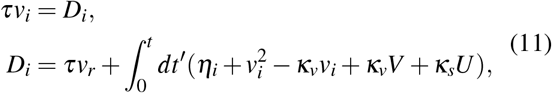

where ∀*i* = 1 · · · *N* and *D* is a functional of the variables *v*_*i*_, 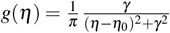 and *U*. We use the same notation mentioned at the beginning of the section 2.1.1.

To couple a variable neuronal density to the Eqs. (11), we can understand neuronal response as the composition over the two possible states. The first one represents the neuron being alive and the second being dead. When the neuron is alive it is expected that its membrane potential satisfies the Eqs. (11) and when it is dead, its membrane potential will be equal to a constant *v*_0_. Now by the definition in subsection 2.1.2, inside *ℬ*_*ε*_ (**x**), the probability of finding the neuron in the alive state is *λ* (**x**, *t*) and consequently the probability of finding the neuron in the death state is 1 − *λ* (**x**, *t*). Then the voltage of the neuron *i* would be the average voltage over the possible states, from which we obtain

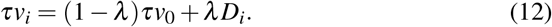

From the Eqs. (12, 11, 7) we can write the microscopical description of neural activity during a neurodegenerative process, in this way we can model the QIF update with a variable density of neurons within *ℬ*_*ε*_ (**x**) as (Appendix 5.3)

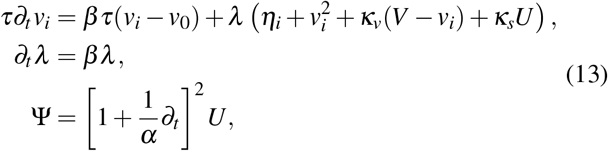

where *i* = 1, · · ·, *N*. As we can see from the Eqs. (13), when we are not in the presence of a neurodegenerative process (*λ* = 1 and *β* = 0), the original QIF model Eqs. (3) and the update QIF model coincide, we can also observe that when in a neighborhood all the neurons have died the potential *V* = *v*_0_, all this mean that Eqs. (13) constitutes a consistent microscopic description with the nature of neurodegenerative processes.

The next step consists of obtaining the macroscopic description of the updated QIF model Eqs. (13) using similar arguments to those developed in Montbrío *et al*. [MPR15]. At the continuous thermodynamic limit (*N* → +∞, *ε* → 0) we can deduce the neural field equation with a variable neuronal density for all **x** ∈ Ω (Appendix 5.4)

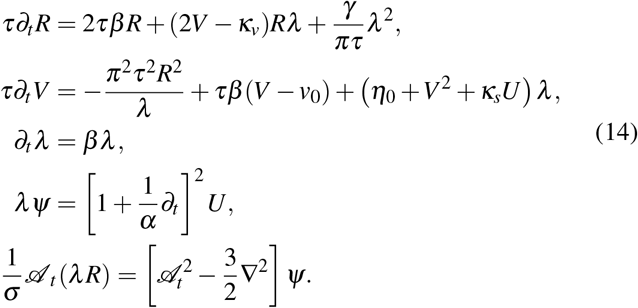

## 3. Results

### 3.1 Theoretical result

The model was stated by Eqs. (14) constitutes our description for neuronal activity during NDDs and our main theoretical result. It is a version of the Next Generation Neural Field models, where we have coupled a variable density of neurons to the field modeled by *λ*. We can also appreciate that the macroscopic state of neuronal activity is described from 5 thermodynamic variables (*R,V,U, ψ, λ*). In the absence of a neurodegenerative processes within the Ω region, we must take *β* (**x**, *t*) = 0 and *λ* (**x**, *t*) = 1 which transforms our model Eqs. (14) into the classical NGNF model Eqs. (4) as expected. This means that our model recovers the model of neuronal activity in the absence of NDDs.

Following the work of Montbrió *et al*. [MPR15], for QIF neuron the Kuramoto order parameter it obtained from the Montbrió parameter *W* which is defined in Appendix 5.4. As for non-neurodegenerative processes, the relationship between the Monbrió parameter and the Kuramoto order parameter holds [MPR15, BRNC22]

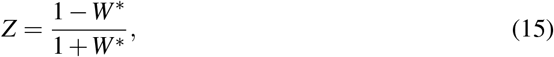

the difference for neurodegenerative processes results from the relationship between the Montbrió parameter *W* and the macroscopic state (*R,V,U, ψ, λ*). As we demonstrate in Appendix 5.4 for our model the exact relation is determined by

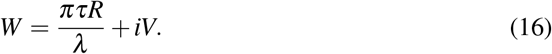

Using Eq. (16) in Eq. (15) we can describe the relationship between the Kuramoto order parameter and the macroscopic state by

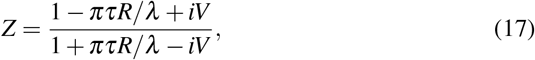

from where we can compute the local synchronization *ζ*_x_(*t*) = | *Z*(x,*t*) | for all **x** ∈ Ω (Appendix 5.5)

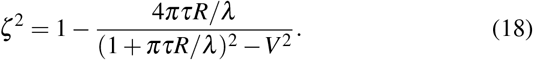

At points **x** ∈ Ω where neurodegeneration is very strong, we have that *λ* (**x**, *t*) → 0, then from expression (18) and the model in Eqs. (14) we can conclude that (Appendix 5.5)

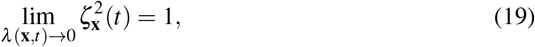

which means that neuronal death induces local synchronization. Although it seems contradictory, this results just reflects the fact that when many neurons are dead, the level of macroscopy activity is very small and with little variation, so, the macroscopic synchronization between two of these areas will appear high as they are almost always in the same state. In the limit, two areas where there are no neurons, will have the same activity (zero) across time and synchronization will be maximum (one).

For the study of more complex synchronization patterns between *K* points within Ω we propose the use of the *K*-points the synchronization, and we will denote as 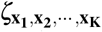. The *K*-point synchronization is an extension of the *ζ* Local synchronization and is calculated from the Kuramoto order parameter of each *K* point as

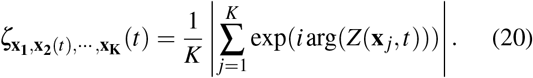

In general, the K-point synchronization is a normalized measure that takes values from 0 (non-synchronized K points) to 1 (for full synchronization). The result expressed in Eq. (19) extends to the K-point synchronization (Appendix 5.6)

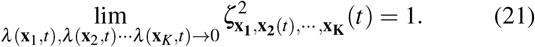

From definition (20) we can compute the two - point synchronization *ζ*_*ab*_ and global synchronization of al neurons in Ω denoted as *ζ*_Ω_. These quantities are of interest because the synchronization of two points resembles the correlation of the firing rate between two points of the neuronal field and the global synchronization offers us a measure of how synchronously the neurons of the entire field work at a given moment. From definition (20) we can show that (Appendix 5.7)

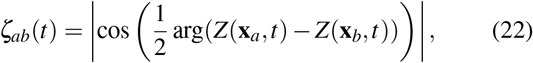

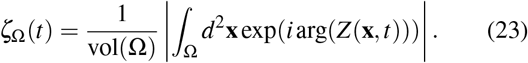

Two-point and global synchronization will be our main tools to study changes in synchronization induced by different settings of neurodegeneration in subsection 3.2.

### 3.2 Experimental results from model simulation

To perform all the simulations of Eqs. (14) we select Ω as a circumference of radius *R* = 8, within which we assume spherical symmetry. As initial values of the model we use a perturbation of the steady state. We only perform the simulation when *β*(**x**, *t*) = 0, *λ*(**x**, *t*) = *λ*_0_ for all **x** ∈ Ω, this scenario represents the case where the level of neurodegeneration varies very slowly and can be considered constant. Other neurodegeneration settings fall outside the objectives of this article.

Fig. 2 show the values of the macroscopic variables of interest *R, V* and *Z* in a radio versus time graph for distinct levels of constant in time and uniform in space neurodegeneration (*λ* (**x**, *t*) = *λ*_0_, *β* (**x**, *t*) = 0). As we can see, for a weak neurodegeneration (*λ* = 0.9) the firing rate (*R*) and the local field potential (*V*) at some moments take values higher than 15 and 200 respectively. As we increase the neurodegeneration until we reach a strong neurodegeneration (*λ* = 0.3), we can see that the amplitude of the variation of the firing rate (*R*) and the local field potential (*V*) decreases, this decrease its appreciable for a strong neurodegeneration where at some moments *R* and *V* takes values not much higher than 0.016 and 1.5, respectively. We can also see how the frequency of spike firing decreases as we increase neurodegeneration. These reflects the fact that increasing neurodegeneration decreases neuronal activity and its intensity, becoming insignificant when neurodegeneration is absolute (*λ* = 0).

**Figure 2.**
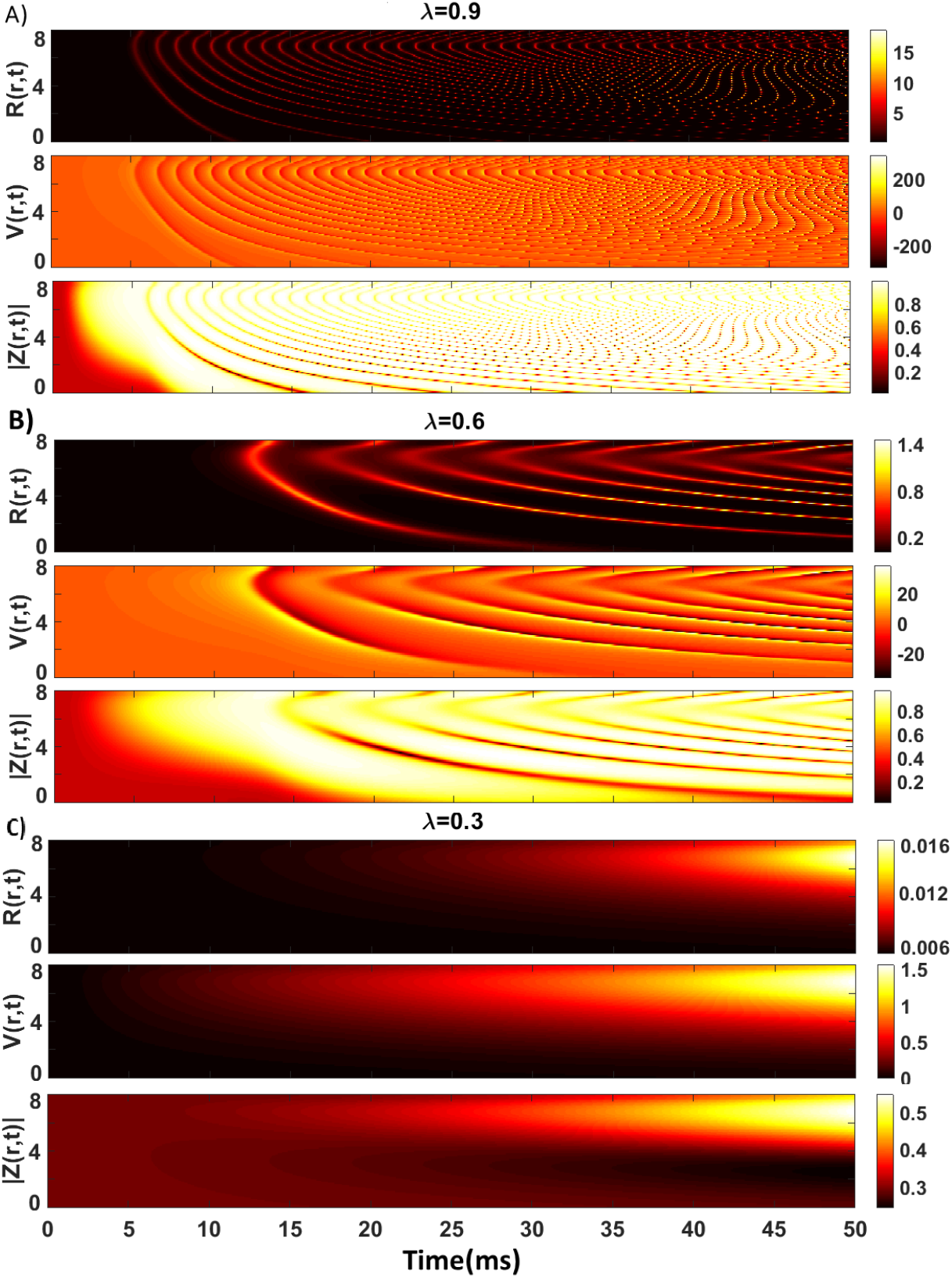
Solutions of the NGNF model with variable neuron density for different levels of neurodegeneration. **A)** Weak neurodegeneration *λ* = 0.9. **B)** Moderate neurodegeneration *λ* = 0.6. **C)** Strong neurodegeneration *λ* = 0.3. Values of the other constants: *τ* = 10, *α* = 3, *ν* = 10, *β* = 0, *v*_0_ = 0, *κ*_*v*_ = 0.9, *κ*_*s*_ = 5, *γ* = 0.5, *η*_0_ = 0.3, *σ* = 1.

Fig. 3 show a space vs space graph of the two - point synchronization *ζ*_*ab*_ for different moments in time and degrees of neurodegeneration. Mathematically this synchronization resembles the functional synchronization between two areas. Varying the degree of neurodegeneration changes the pattern in two - points synchronization for all instant in time. In the last column of Fig. 3, we can appreciate the mean value of the two - points synchronization ⟨*ζ*_*ab*_⟩ in time for different degrees of neurodegeneration. In it we note a marked difference between the case of moderate and strong neurodegeneration both in spatial distribution and in intensity. In general, the maximum of ⟨*ζ*_*ab*_⟩ tend to decrease for *λ* = 0.9 to *λ* = 0.3, while for values smaller than 0.3 the maximum of ⟨*ζ*_*ab*_⟩ starts to increase reaching ⟨*ζ*_*ab*_⟩ = 1 for all points when neurodegeneration is absolute.

**Figure 3.**
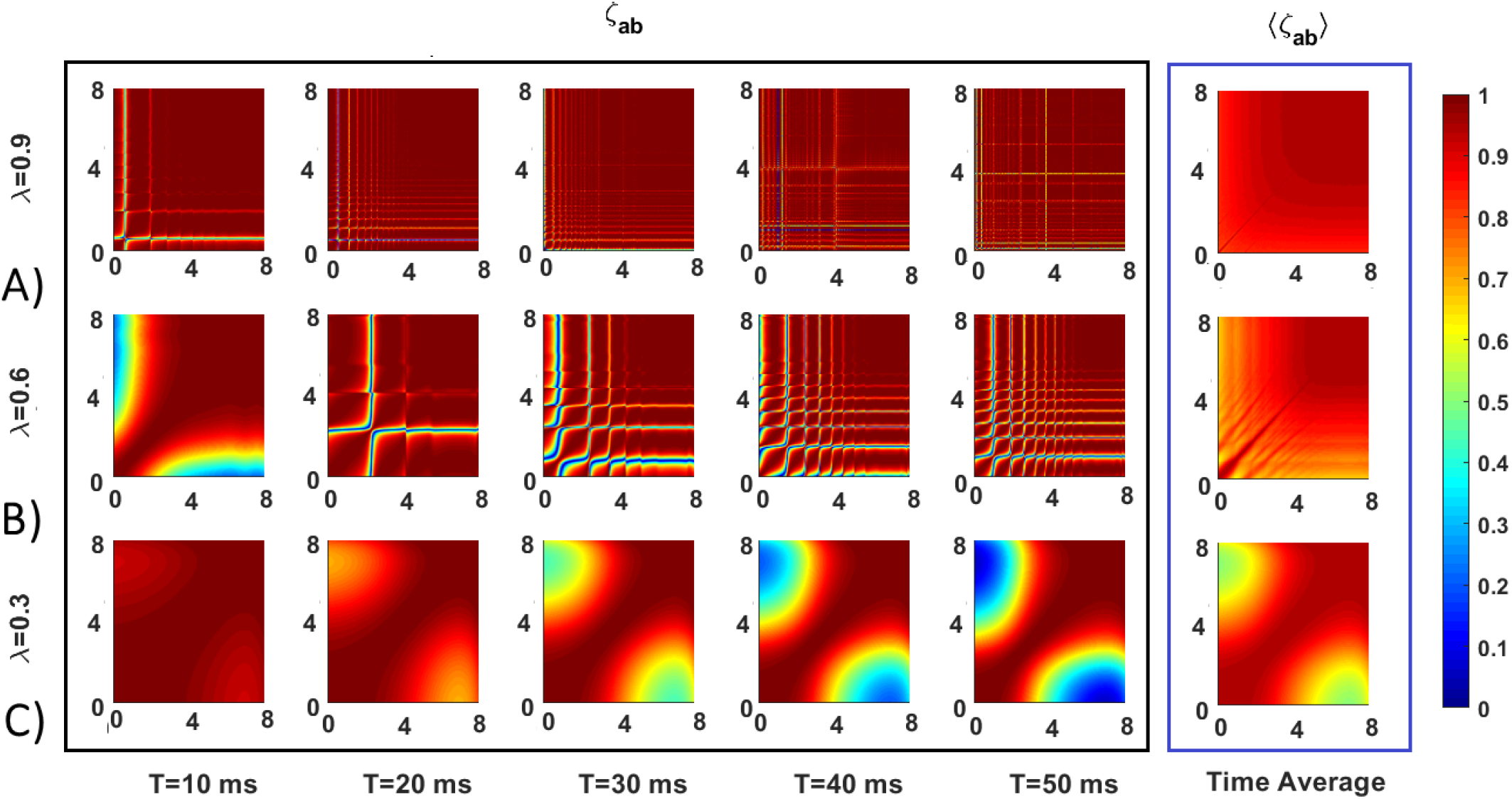
Two point synchronization over time for different level of neurodegeneration. **A)** Weak neurodegeneration (*λ* = 0.9). **B)** Moderate neurodegeneration (*λ* = 0.6). **C)** Strong neurodegeneration (*λ* = 0.3).Values of the other constants: *τ* = 10, *α* = 3, *ν* = 10, *β* = 0, *v*_0_ = 0, *κ*_*v*_ = 0.9, *κ*_*s*_ = 5, *η*_0_ = 0.3, *γ* = 0.5, *σ* = 1.

Finally, Fig. 4 show the evolution in time of the global synchronization inside Ω for different level of neurodegeneration. As can observe, global synchronization tend to decrease their mean value as we increase neurodegneration form *λ* = 0.9 to *λ* = 0.3. In particular for strong neurodegeneration (*λ <* 0.3) we can appreciate a hypersynchronization of higher magnitude that for weak neurodegeneration.

**Figure 4.**
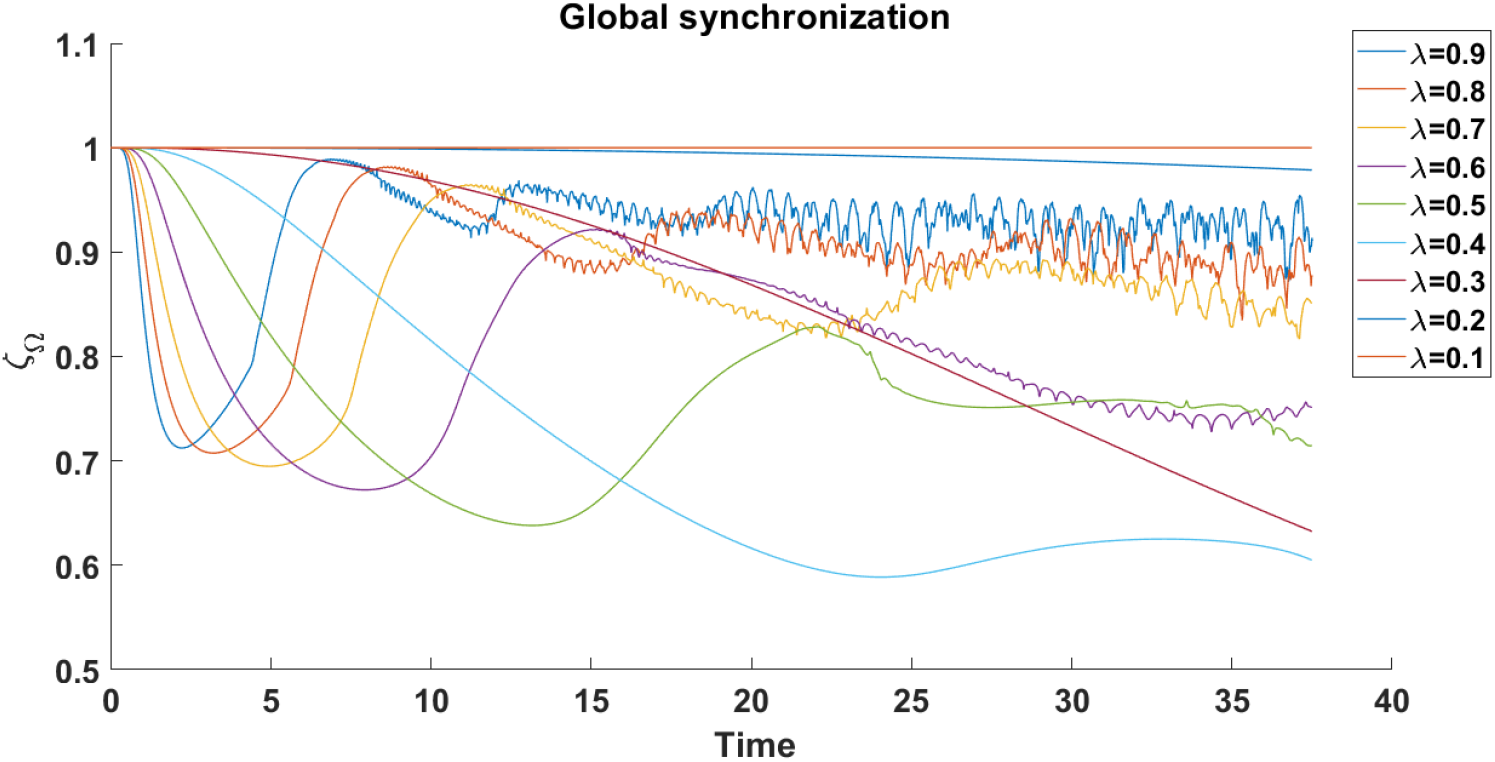
Global synchronization for different level of neurodegeneration. Values of the other constants: *τ* = 10, *α* = 3, *ν* = 10, *v*_0_ = 0, *κ*_*v*_ = 0.9, *κ*_*s*_ = 5, *η*_0_ = 0.3, *γ* = 0.5, *σ* = 1.

## 4. Discussion

In this work we have obtained NGNF model with a variable density of neurons (section 2), which allows us to study neural activity during neurodegenerative processes. To do this, we considered a series of hypotheses about the microscopic nature of neurodegeneration and its relationship with the electrical proprieties of neurons (subsection 2.1). From the microscopic description by an updated QIF model Eqs. (13) we find the exact macroscopic description of neural activity Eqs. (14) in the presence of neurodegenerative processes (subsection 2.2). As a result, the description of the macroscopic state of neural activity is determined from 5 variables (subsection 3.1), which means that when modeling neurodegenerative processes, we are adding degrees of freedom to the NGNF model.

The hypotheses of our model were chosen in a consistent manner with the nature of neurodegenerative processes (subsection 2.1), however, these hypotheses constitute a simplification of the reality of neurodegeneration, which may present limits in their application for different NDDs. To validate our model, we have only evaluated the consistency of the simulation results (subsection 3.2) with results from the literature exposed in the Introduction 1. This means that we do not currently have direct experimental data to prove the quantitative validity of the developed model. In general, when studying a particular NDD we could add stronger hypotheses to the developed model to guarantee a better precision of the predictions.

As we could see during the presentation of the theoretical results (subsection 3.1), under the hypotheses assumed the relationship between the macroscopic state (*R,V,U, ψ, λ*) and the Kuramoto order parameter Z, differs from previous results for non-neurodegenerative processes [MPR15, BRNC22]. We observe that for NDDs, for the direct calculation of the local synchronization or Kuramoto order parameter, it is necessary to consider the degree of neurodegeneration determined by *λ* using the expression

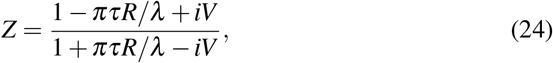

this since neurodegeneration influences directly the synchronization patterns. Using this relationship we were able to demonstrate analytically that an absolute neurodegeneration at one point induces a local hypersynchronization at that point.

We also observed that increasing the degree of neurodegeneration in our model decreases neural activity and its intensity. Result that is consistent with experimental observation, since if the neurons within a region die due to some NDDs, we should observe less neural activity. On the other hand, we were able to verify that the increase in neurodegeneration produces changes in the two - point synchronization *ζ*_*ab*_ a measure of the synchronization between points of the Neural Field. During the subsection 3.2 we observe how two points of the Neural Field are synchronize and desynchronize, the temporal average for different neurodegeneration degrees of this process ⟨*ζ*_*ab*_⟩ translates into a differentiated correlated activity between spatially separated areas. In the literature it is common to measure the correlation between neuronal activity of different brain areas through functional connectivity [FFF^+^96, LBK^+^10, UCKB^+^09]. Previous research use functional connectivity as a feature to classify and measure the progress of NDDs [PFVDH^+^14b, SDHD^+^09, SJN^+^07, SBD^+^08, WLW^+^11]. If we take into account the well - known hypothesis that long - range functional connectivity between brain areas is driven by synchronization mechanisms [EGHN13, Fri97, PADH^+^15], the results of this work could indicate that the reason why functional connectivity is a good feature of NDDs is due to the direct relationship between synchronization and neuronal density expressed in the average two - point synchronization ⟨*ζ*_*ab*_⟩.

Finally, we observed changes in the global synchronization *ζ*_Ω_ associated with neurodegenerative processes. Particularly in presence of a strong neurodegeneration (*λ <* 0.3) a *ζ*_Ω_ ∼ 1 hyper synchronization is observed, which is a consistent result, since if all neurons within a given region have died, they are all synchronized in the same state. On the other hand, we appreciate that gradually increasing from a weak to moderate neurodegeneration (0.3 *< λ <* 0.9), produces a marked decrease in global synchronization. In the literature it is considered that neuronal synchronization has an important role in the execution of cognitive processes such as memory, movement etc [BD04, BLP14, TB03, Wan10, CA05a]. The results obtained in this work indicate that the progress of neurodegeneration is related to neuronal synchronization, both local and spatial, which could mean an alteration of cognitive processes i.e. the neurological symptoms associated with the different NDDs such as akinesis, memory impairment, tremor etc.

During future research we will study if the results obtained are maintained for other combinations of the parameters of the model. We will study other more dynamic neuronal neurodegeneration configurations, such as the one with *β <* 0. This type of configuration, for large values of |*β*| could be especially useful in the study of aggressive neurodegeneration as in the case of Glioblastomas in the exponential phase. It would also be ideal to carry out laboratory experiments that would allow us to validate our results and, if necessary, consider other more specific hypotheses as part of the model design.

## 5. Appendix

### 5.1 Dynamics of *λ*

If *ρ*_*H*_ is the health neuronal density inside Ω. Then by the definition of *λ*_*ε*_ (**x**, *t*) (subsection 2.1.2)

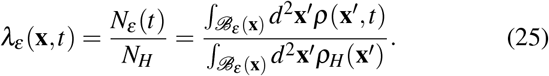

Using the Mean Value Theorem of the Integral Calculus, is possible to reduce Eq. (25) as

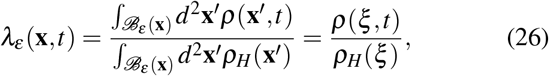

where *ξ* belongs to *ℬ*_*ε*_ (**x**). Taking the continuum limit (*ε* → 0) result

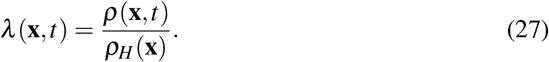

Finally it is possible to deduce Eq. (7) by substituting Eq. (27) into Eq. (5) and taking *λ*_0_ = *ρ*_0_*/ρ*_*H*_.

### 5.2 Brain Wave Equation

By definition (see Subsection 10), the conditional spatio - temporal drive *ψ* is

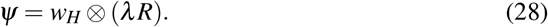

Expanding the expression we cant represent Eq. (28) as a convolution product as

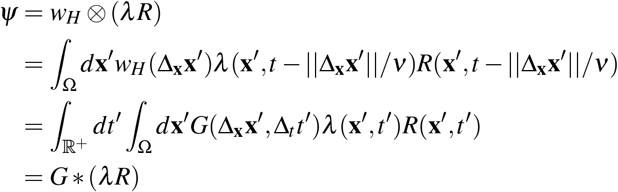

where *G*(*x, t*) = *w*_*H*_(*x*)*δ* (*t* − *x/ν*) and ∗ is the spatio - temporal convolution. Now form the convolution representation of the conditional spatio - temporal drive *ψ* we cant derive a PDE model, for that we have to takes Fourier Transformation^6^ in both sides of the convolution expression of *ψ*

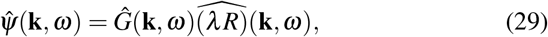

where

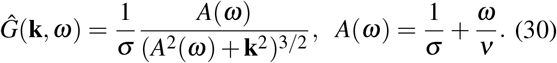

Then from Eq. (30) we have

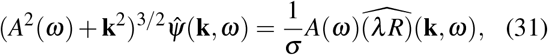

where taking the inverse Fourier Transform and the long wave approximation (||**k**|| ∼ 0) result

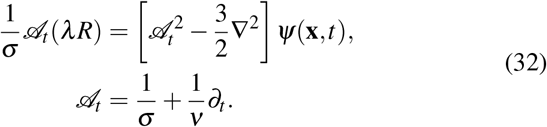

### 5.3 Differential form of the update QIF model

To obtain the differential form of the update QIF model we have to differentiate the Eq. (12) taking into account Eq. (7) as

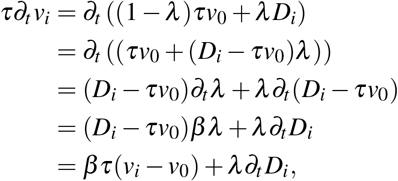

where differentiating the functional operator *D*_*i*_ with respect to time we conclude Eq. (13).

### 5.4 Thermodynamic limit of the update QIF model

To obtain the macroscopic description of neural activity, first we must find the thermodynamic limit (*N* → ∞) of the Eqs. (13) inside *ℬ*_*ε*_ (**x**) ⊂ Ω, and then express the continuum limit (*ε* → 0) for all **x** ∈ Ω. At the thermodynamic limit we can write the continuity equation of the voltage distribution *F*(*v*|*η*, **x**, *t*) in ℬ_*ε*_ (**x**) ⊂ Ω

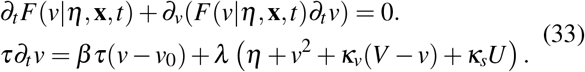

Following the work in [MPR15, BRNC22, BOF^+^20], we can solve the Eq. (33) by applying the LA, which consists of applying the following formula on Eq. (33)

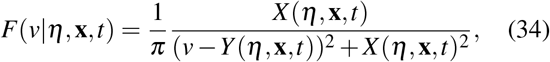

For a fixed value of *η* we can find the firing rate *r*(*η*, **x**, *t*) as the probability flux at infinity *F*(*v* → ∞, **x**, *t*)*∂*_*t*_*v*(*v* → ∞). Using the LA and taking the limit to infinity we have

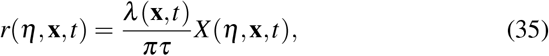

where averaging over the distribution of *η* we obtain a formula for the firing rate

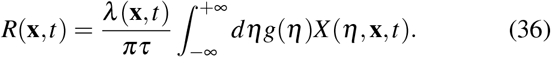

On the other hand it is possible to establish for *Y* (*η*, **x**, *t*) the next formula by contour integration

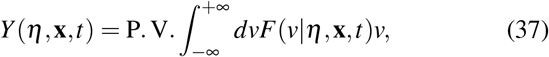

where averaging over the distribution of *η* as before, we obtain a formula for the average potential

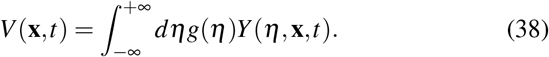

We can obtain an equation for *W*_*η*_ = *X* (*η*, **x**, *t*) + *iY* (*η*, **x**, *t*) By the substitution of Eq. (34) into the continuity equation and balancing powers of *v* as

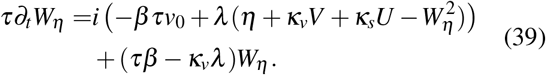

From Eq. (39) we can find a equation for the average of *W*_*η*_ over the distribution of *η* denoted by *W*. Using the Lorentzian distribution *g*(*η*) = (*γ*/π)/((*η* − *η*_0_)^2^ + *γ* ^2^) we have for *W*

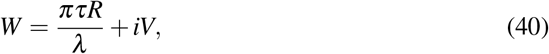

and from Eq. (39)

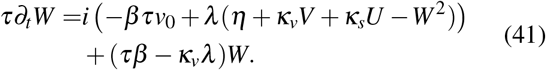

In the literature the variable *W* is called the Montbrió parameter and from it the Kuramoto order parameter is determined.

Substituting directly the formula (40) in Eq. (41) we obtain a equation for the firing rate *R* and the average potential *V* inside *ℬ*_*ε*_ (**x**) ⊂ Ω by

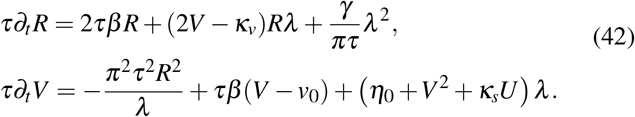

Finally we can find the continuum limit of the Eqs. (42) to obtain the Neural Field model with a variable neuronal density for all **x** ∈ Ω

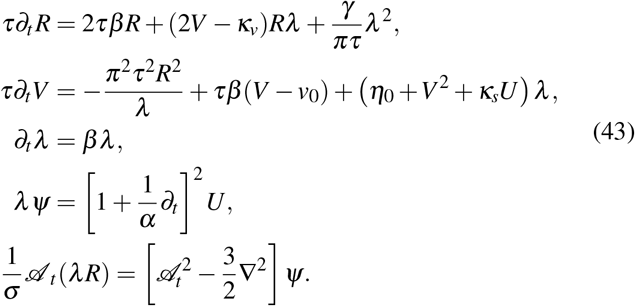

### 5.5 Local syncronization

The local synchronization by definition is the modulus of the Kuramoto order parameter for the given point *ζ* ^*2*^ = |*Z*|^2^, expanding this expression we get

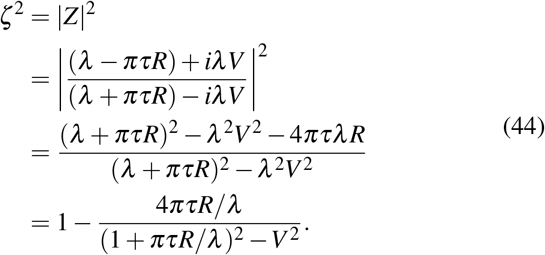

To analyze the behavior of *ζ*^2^ when all the neurons died or *λ* → 0, it is enough to analyze the behavior of the firing rate. From our mathematical model Eqs. (14) when *λ* → 0 we can neglect the terms that depend on *λ* in the equation of the firing rate and analyze instead the equation

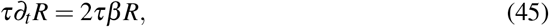

this equation have for solution *R* = *R*_0_ exp(2*τβt*), where using the *λ* equation Eq. (7) reduces to

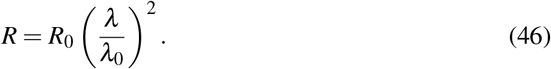

Substituting the expression (46) in Eq. (44) when *λ* → 0 we have

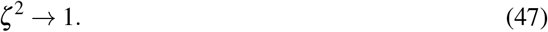

### 5.6 K - point synchronization

It is clear for the model hypothesis that

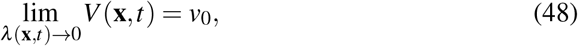

also from expression (46)

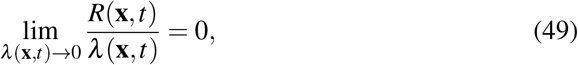

then from expressions (48), (49) and the expression of the Kuramoto order parameter for neurodegenerative processes (17) we have

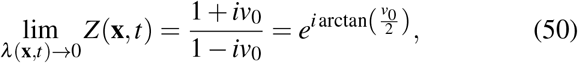

then in general

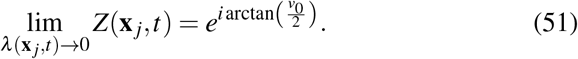

Using the definition of the *K* point synchronization and expression (51)

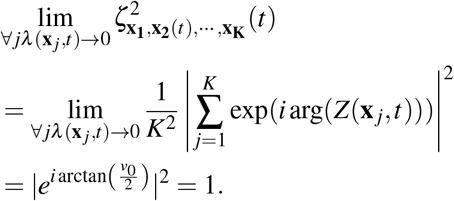

### 5.7 Two - point synchronization formula

From the definition of two - point synchronization

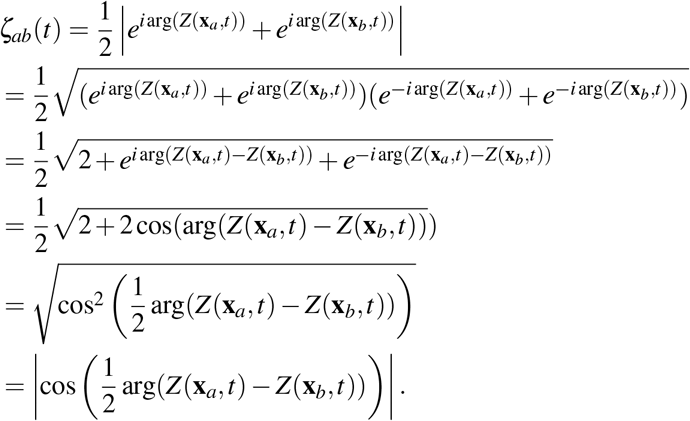

## Footnotes

1 A symptom of PD associated with inability to initiate movement and slowness of movement

2 Local synchronization at a point **x** refers to the synchronization of neurons in a small neighborhood of **x**

3 *ℬ*_*ε*_ (**x**) is a ball with center at **x** ∈ Ω, radius *ε* and Ω = ∪_**x**∈Ω_ *ℬ*_*ε*_ (**x**)

4 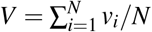.

5 (*f* ⊗ *g*)(**x**, *t*) = ∫_Ω_ *d*^2^**x**′ *f*(**x, x**′)*g*(**x**′, *t* − ‖Δ_**x**_ **x**′‖/*v*)

6 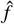 denote de direct Fourier Transform

